# Processing of Emotional Faces Has Unique Functional and Cytoarchitectural Associations in those with an Autism Spectrum Diagnosis

**DOI:** 10.1101/2025.11.13.688249

**Authors:** Zachary P. Christensen, Edward G. Freedman, John J. Foxe

**Affiliations:** The Frederick J. and Marion A. Schindler Cognitive Neurophysiology Laboratory, The Del Monte Institute for Neuroscience Department of Neuroscience, University of Rochester School of Medicine and Dentistry, Rochester, New York 14642, USA; The Golisano Intellectual and Developmental Disabilities Institute, University of Rochester School of Medicine and Dentistry, Rochester, New York 14642, USA

## Abstract

Those with an autism spectrum diagnosis (ASD) have been found to process emotional faces differently than other populations. Processing of emotional faces requires engagement of temporal, frontal, occipital, and limbic brain regions. Functional activity has been shown to differ in those with an ASD, particularly in the amygdala, inferior frontal cortex (IFC), and temporal brain regions. However, the consistency and direction of these associations have been inconsistent across studies. Recent findings have demonstrated that measures of neuron density differ in those with an ASD. Some of these regional differences in cytoarchitecture coincide with regions important to emotional facial processing. Therefore, the interaction between cytoarchitecture and functional activity may be important in elucidating unique neurophysiology in ASD. The present study uses diffusion weighted imaging (DWI) and functional magnetic resonance imaging (fMRI) data from the Adolescent Brain Cognitive Development^sm^ (ABCD®) study to investigate the relationship between cytoarchitecture and functional activity during emotional face processing in those with an ASD. 75 individuals with a reported ASD and 6,396 individuals with no reported diagnosis of an ASD (nASD) were identified. The emotional n-back (EN-Back) task was administered during fMRI acquisition, activating regions of the brain associated with processing emotional faces. Neuron cell body density was positively correlated with functional activation in the left amygdala in the ASD group but not the nASD group. These findings suggest that a unique relationship may exist between neuron cell body density in the left amygdala and functional activity while processing emotional faces in those with an ASD.

## 1 Introduction

Differences in social communication and social interactions are among the defining characteristics in individuals who receive an autism spectrum diagnosis (ASD) (American Psychiatric Association, 2013). Communicating emotions through facial expressions is critical to social interactions, and people with an ASD have been found to process emotional faces differently than other populations (Griffin et al., 2021; Uljarevic & Hamilton, 2013; Vanneau, 2025). The behavioral presentation of abnormal facial processing in autism includes poor memory (Boucher et al, 1998), differences in eye-tracking (Sterling et al., 2008), deviations in perception of emotion (Ashwin et al., 2006), and difficulty evaluating trustworthiness (Adolphs et al., 2001). Despite the well-established association of ASD and atypical facial processing, the underlying functional-structural relationships potentially driving these behavioral manifestations are still poorly understood (Webb et al., 2011).

Human processing of faces encompasses a variety of steps that differentially engage neuroanatomical regions depending on task-specific demands (Kanwisher & Barton, 2011). The occipital face area and fusiform facilitate processing of visual stimuli related to the face (Kanwisher et al., 1997; Kanwisher & Barton, 2011; Kietzmann et al., 2015). The fusiform plays a particularly important role in simple face detection (Solomon-Harris et al., 2013), and the amygdala is primarily involved with discrimination between emotional and neutral faces (Sato et al., 2021; Vuilleumier et al., 2004). Further higher-order processing of emotional faces engages the inferior frontal cortex (IFC) (Elbich & Scherf, 2017; Rämä & Courtney, 2005), and the uncinate fasciculus which connects regions in the temporal lobe (Coad et al., 2020). While clear and distinct tasks have helped elucidate key regions in the processing of emotional faces, deficits related to psychiatric and developmental disorders such as ASD appear to produce more subtle changes in neurophysiology.

Previous investigations found that simple processing of emotional faces in ASD was associated with greater functional activity in the amygdala (Monk et al., 2010) and decreased functional connectivity in the inferior frontal regions in the context of working memory tasks (Rämä & Courtney, 2005). In contrast, more recent investigations have found decreased functional activation in the fusiform gyrus (Nickl-Jockschat et al., 2015) and decreased functional activity and connectivity in the amygdala (Guo et al., 2016; Kleinhans et al., 2011) in relation to emotional faces. These discrepancies may be a result of overly general measures obscuring more discrete changes. Functional activity in the amygdala has been reported to be more distinctly related to fear than other emotional stimuli in ASD (McKechanie et al., 2022). Other studies have identified altered habituation of the amygdala to emotional stimuli in ASD (Tam et al., 2017). However, seeking novel approaches to characterize these neuroanatomical regions permits leverage of novel techniques without limiting the scope of behavioral responses being considered.

Recent investigation of the fusiform found that predictive models trained on features derived from multiple modalities of structural and functional imaging, better predicted social outcomes in individuals with an ASD than models trained on features of a single modality (Floris et al., 2025). Although models reducing many features derived from multiple modalities are difficult to interpret, their predictive accuracy may indicate that properly characterizing the relationship between neurophysiology and social outcomes requires multiple modalities. For example, administering a selective-serotonin reuptake inhibitor to those with an ASD shifts functional activity of serotonergic neurons in the ventromedial prefrontal cortex towards neurotypical individuals when viewing emotional faces (Wong et al., 2022). Therefore, changes in functional activity may be tied to characteristics of specific neuron populations. Differences in neuron populations of those with ASD have been found in non-task associated analyses of cytoarchitecture (Christensen et al., 2024). Individuals with ASD have been found with a reduced number of amygdalar neurons (Schumann & Amaral, 2006), fewer cerebral Purkinje cells (Kemper & Bauman, 1998), smaller Purkinje cells in the cerebellum (Fatemi et al., 2002), patches of cortical disorganization (Stoner et al., 2014), and a greater number of narrow minicolumns in the frontal and temporal lobes (Buxhoeveden et al., 2006; M. F. Casanova et al., 2002, 2002; M. Casanova & Trippe, 2009). However, relatively little is known about emotional facial processing and cytoarchitecture in ASD.

The present study sought to identify unique associations between functional activity and neuron density, building on previous findings in the Adolescent Brain and Cognitive Development ^sm^ (ABCD®) study (Barch et al., 2018, 2021; Casey et al., 2018; Hagler et al., 2019). In addition to selecting regions implicated in functional processing of emotional faces, the present study selected regional and cytoarchitectural measures based on notable differences that were previously detected in those with an ASD in the ABCD study cohort. Specifically, the present study focused on recent findings that demonstrated those with an ASD had increased restricted normalized directional diffusion (RND) within the inferior frontal and fusiform cortices, and increased restricted normalized isotropic diffusion (RNI) within the amygdala (Christensen et al., 2024). Our hypothesis was that the magnitude of these changes within individuals with ASD would correspond to changes in functional activity while processing emotional faces, when contrasted to those without an ASD (nASD).

## 2 Methods

### 2.1 Study Population

Data from the Adolescent Brain and Cognitive Development^sm^ (ABCD®) study 6.0 release were used (Jernigan et al., 2025). The ABCD study consists of 11,873 children recruited between ages 9-10-year-old at baseline. Participants were recruited from 22 sites (21 active sites) across the US from public and private schools and selection was based on race and ethnicity, sex, socioeconomic status, and urbanicity, in an effort to match US population and demographics (Garavan et al., 2018). The ABCD study is currently in the process of following these individuals over the course of 10 years - https://abcdstudy.org/. The present study utilized imaging data collected at the ABCD study’s baseline, 2-year follow-up, 4-year follow-up, and 6-year follow-up. Parent consent and child assent were approved by the central Institutional Review Board at the University of California – San Diego. Study procedures were conducted in accordance with the tenets of the declaration of Helsinki.

Individuals were excluded if they were not able to complete all administered tasks or complete an MRI scan. This excluded individuals who lacked fluency in English or uncorrectable sensory deficits (e.g., legal blindness). Participants were also excluded if they had a psychiatric diagnosis that prevented attendance at regular classes in school, neurological issues (e.g., brain tumor, previous head injury with loss of consciousness > 30 minutes), or significant perinatal medical issues (e.g., gestational age < 28 weeks, birthweight <1.2 kg, complications resulting in >1 month of hospitalization following birth) (Garavan et al., 2018). Individuals were excluded if they did not pass quality control measures, which included both automated and manual methods such as findings of artifact or file corruption by a radiologist (Chaarani et al., 2021). Given the focus of the current investigation, individuals that were missing pertinent imaging data (see Neuroimaging Data section) or had a task accuracy below 60% (see Emotional N-Back section) were excluded. Representation of demographics used in the present study are presented in Table 1.

**Table 1.**
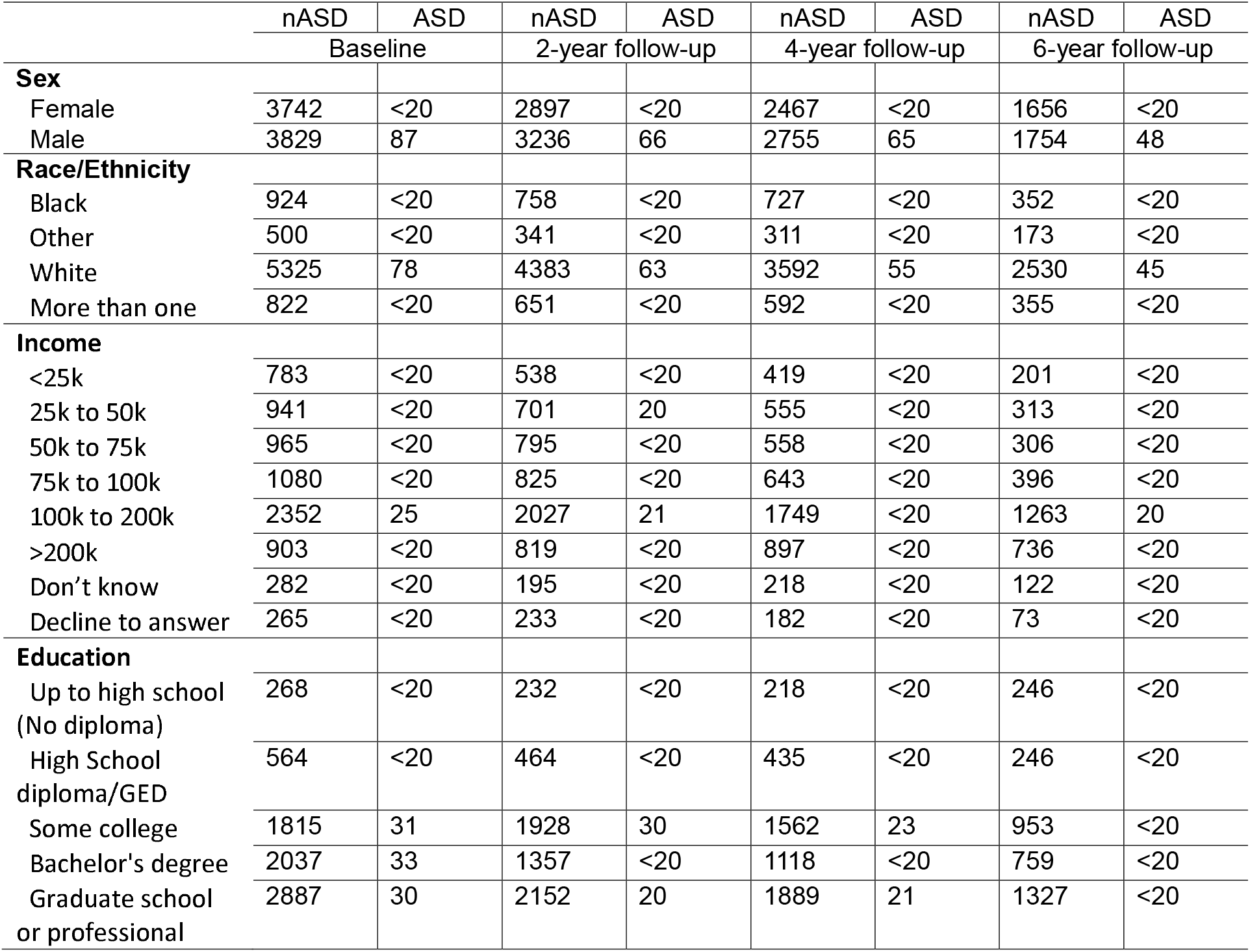
Demographics for the nASD and ASD groups across all timepoints where imaging was collected. Exact values are only reported where a subgrouping contained 20 or more individuals to reduce risk of identifying individuals.

|  | nASD | ASD | nASD | ASD | nASD | ASD | nASD | ASD |
| --- | --- | --- | --- | --- | --- | --- | --- | --- |
|  | Baseline |  | 2-year follow-up |  | 4-year follow-up |  | 6-year follow-up |  |
| **Sex** |  |  |  |  |  |  |  |  |
| Female | 3742 | <20 | 2897 | <20 | 2467 | <20 | 1656 | <20 |
| Male | 3829 | 87 | 3236 | 66 | 2755 | 65 | 1754 | 48 |
| **Race/Ethnicity** |  |  |  |  |  |  |  |  |
| Black | 924 | <20 | 758 | <20 | 727 | <20 | 352 | <20 |
| Other | 500 | <20 | 341 | <20 | 311 | <20 | 173 | <20 |
| White | 5325 | 78 | 4383 | 63 | 3592 | 55 | 2530 | 45 |
| More than one | 822 | <20 | 651 | <20 | 592 | <20 | 355 | <20 |
| **Income** |  |  |  |  |  |  |  |  |
| <25k | 783 | <20 | 538 | <20 | 419 | <20 | 201 | <20 |
| 25k to 50k | 941 | <20 | 701 | 20 | 555 | <20 | 313 | <20 |
| 50k to 75k | 965 | <20 | 795 | <20 | 558 | <20 | 306 | <20 |
| 75k to 100k | 1080 | <20 | 825 | <20 | 643 | <20 | 396 | <20 |
| 100k to 200k | 2352 | 25 | 2027 | 21 | 1749 | <20 | 1263 | 20 |
| >200k | 903 | <20 | 819 | <20 | 897 | <20 | 736 | <20 |
| Don't know | 282 | <20 | 195 | <20 | 218 | <20 | 122 | <20 |
| Decline to answer | 265 | <20 | 233 | <20 | 182 | <20 | 73 | <20 |
| **Education** |  |  |  |  |  |  |  |  |
| Up to high school<br>(No diploma) | 268 | <20 | 232 | <20 | 218 | <20 | 246 | <20 |
| High School<br>diploma/GED | 564 | <20 | 464 | <20 | 435 | <20 | 246 | <20 |
| Some college | 1815 | 31 | 1928 | 30 | 1562 | 23 | 953 | <20 |
| Bachelor's degree | 2037 | 33 | 1357 | <20 | 1118 | <20 | 759 | <20 |
| Graduate school<br>or professional | 2887 | 30 | 2152 | 20 | 1889 | 21 | 1327 | <20 |

### 2.4 Diagnostic Groups and Cognitive Assessments

The ABCD cohort was subdivided into those with a presumed diagnosis of ASD and those with no ASD diagnosis (nASD), as previously reported (Christensen et al., 2024). The brief social responsiveness scale (SRS) was used to overcome the absence of formal testing or clinical records to confirm a diagnosis of ASD. A T-score above 60 on the SRS has demonstrated good reliability and sensitivity for identifying ASD across various psychopathologies (Moul et al., 2015). Participants who had a parent reported diagnosis of ASD and scored below this threshold on the SRS were removed from subsequent analysis to improve validity of diagnostic categories, with 164 meeting for ASD at baseline. Those who met criteria for ASD and had the requisite measurers for the present study are presented in Table 1.

### 2.3 Emotional N-Back

The emotional n-back (EN-Back) was used to engage processing of emotional faces across 160 trials (Barch et al., 2013). A series of emotional faces (happy, fearful, neutral) were presented to children. There were two variants of the task where children pressed a button indicating a match for the image immediately preceding a given image (0-back) or the image two iterations ago (2-back). Four task blocks of each variant of the task were administered in approximately 5-minute fMRI runs. Blocks were composed of trials consisting of a 2 second stimulus (image) followed by a 500-millisecond fixation cross. Individual faces were presented with only one unique facial expression at a time. 24 unique happy, fearful, and neutral faces were presented (72 in total). Children were familiarized with the task in a practice session prior to administration of the actual task. Similar to prior work on the N-Back task in the ABCD study, participants with a total accuracy below 60% were excluded (Pedersen et al., 2023). This resulted in dropping 3 individuals in the ASD group and 271 individuals in the nASD group (**see Error! Reference source not found**.).

### 2.4 Neuroimaging Data

In-depth description of image acquisition, processing, and normalization are fully provided elsewhere (Hagler et al., 2019). Image protocols for the present study include 3D T1-weighted (T1w) acquisition (1 mm isotropic) magnetization-prepared rapid acquisition gradient echo scans, 3D T2-weighted (T2w) acquisition (1 mm isotropic) fast spin echo with variable flip angle scans, and diffusion weighted imaging (DWI) acquisition (1.7 mm isotropic) multiband EPI scans with 96 noncollinear gradient directions (6 directions at b = 500 s/mm^2^, 15 directions at b = 1000 s/mm^2^, 15 directions at b = 2000 s/mm^2^, and 60 directions at b = 3000 s/mm^2^) and seven b = 0 s/mm^2^ frames. fMRI acquisition (2.4 mm isotropic, TR=800 ms) employed multiband echo planar imaging (EPI).

Images underwent standard preprocessing as previously described (including corrections for eddy currents, motion, field inhomogeneities as well as other potential sources of artifact) (Casey et al., 2018). The ABCD study protocol also includes quality control measures that facilitated removing data where artifact due to noise could not be appropriately corrected in preprocessing (Hagler et al., 2019). In brief, diffusion weighted images were fit to a restriction spectrum imaging (RSI) model (White et al., 2013). RSI provides multicompartmental measures of diffusion unique to intracellular (restricted) and extracellular (hindered) compartments of brain tissue. TND represents the summed estimates of restricted normalized directional diffusion (RND; corresponding to axonal structures) and restricted normalized isotropic diffusion (RNI; corresponding to the neuron body). The classic DTI model excluded frames greater than 1000 s/mm2, corresponding to the classic single-b-value acquisition sequence. fMRI time-series data was modeled with nuisance regressors such as motion and respiration. Estimates of activation during each fMRI task block were derived using a general linear model (GLM).

Measures of functional activity and neuron density were sampled in the right and left hemisphere for the inferior frontal cortex, fusiform cortex, and amygdala. These regions were selected due to their implication in previous literature for emotional facial processing along with identification in prior investigation of neuron density in the ABCD cohort (Christensen et al., 2024). These measures were derived from cortical and subcortical regions found in the Desikan-Killiany brain atlas (Klein & Tourville, 2012). All measures were pulled from the curated ABCD 6.0 data release.

### 2.5 Statistical Analysis

A mixed effects linear model was used to analyze the relationship between functional activity and neuron density, using covariates such as sex, highest parent education, collection site, reported use of medications for ADHD (methylphenidate, Ritalin, Adderall, etc.). Each of these models was compared to the null model via likelihood ratio tests, as done in previous studies, to avoid over parameterization (Bates et al., 2014; Christensen et al., 2021), using the MixedModels package (Douglas Bates et al., 2019). Benjamini-Hochberg correction was used to correct for multiple comparisons (Benjamini & Hochberg, 1995).

## 3 Results

### 3.2 Functional Activity

The ASD group had lower functional activity in the IFC bilaterally (left: β=−0.102119, SE=0.104189, p-value=0.3270; right: β=−0.024506, SE=0.111295, p-value=0.8257). The ASD group had lower activity in the left fusiform (β=−0.0499036, SE=0.108879, p-value=0.6467) and higher activity in the right fusiform (β=0.039266, SE=0.114924, p-value=0.7326). The ASD group had lower functional activity in the amygdala bilaterally (left: −0.332826, SE=0.106454, p-value=0.0018; right: β=−0.0328482, SE=0.128134, p-value=0.7977). Therefore, the ASD group demonstrated a statistically significant reduction in functional activity while processing emotional faces versus neutral faces.

When compared to females, males had lower functional activity measures in the left IFC (β=−0.00108806, SE=0.00467593, p-value=0.8160) and higher in the right IFC (β=0.000224535, SE=0.00478785, p-value=0.9626). Similarly, males had lower activity in the left fusiform (β=−0.0499036, SE=0.108879, p-value=0.6467) and higher in the right fusiform (β=0.00407826, SE=0.00497943, p-value=0.4128). Males had higher functional activity bilaterally in the amygdala (left: β= 0.00167443, SE=0.00530967, p-value=0.7525; right: β=−0.0328482, SE=0.128134, p-value=0.7977). Altogether, these demonstrated modest relationship between sex and functional activity when processing emotional faces, failing to reach statistical significance.

### 3.2 Neuron Density and Function Activity

Although the linear relationship between left IFC RND and functional activity initially demonstrated a negative correlation (see Figure 1 top-left), inclusion of demographic control variables resulted in a non-significant positive relationship in the ASD group (β=0.13276, SE=0.499362, p-value=0.7903). The initially apparent strong correlation between left IFC RND and functional activity in the ASD group remained positive but was greatly attenuated once accounting for confounding variables (β=0.396676, SE=0.499215, p-value=0.4268). The negative association between right IFC RND and functional activity (see Figure 1 top-right) remained modest across both the nASD group (β= −0.117065, SE=0.529851, p-value=0.8251) and the ASD group (β=−0.0186091, SE=0.529754, p-value=0.9720).

**Figure 1.**
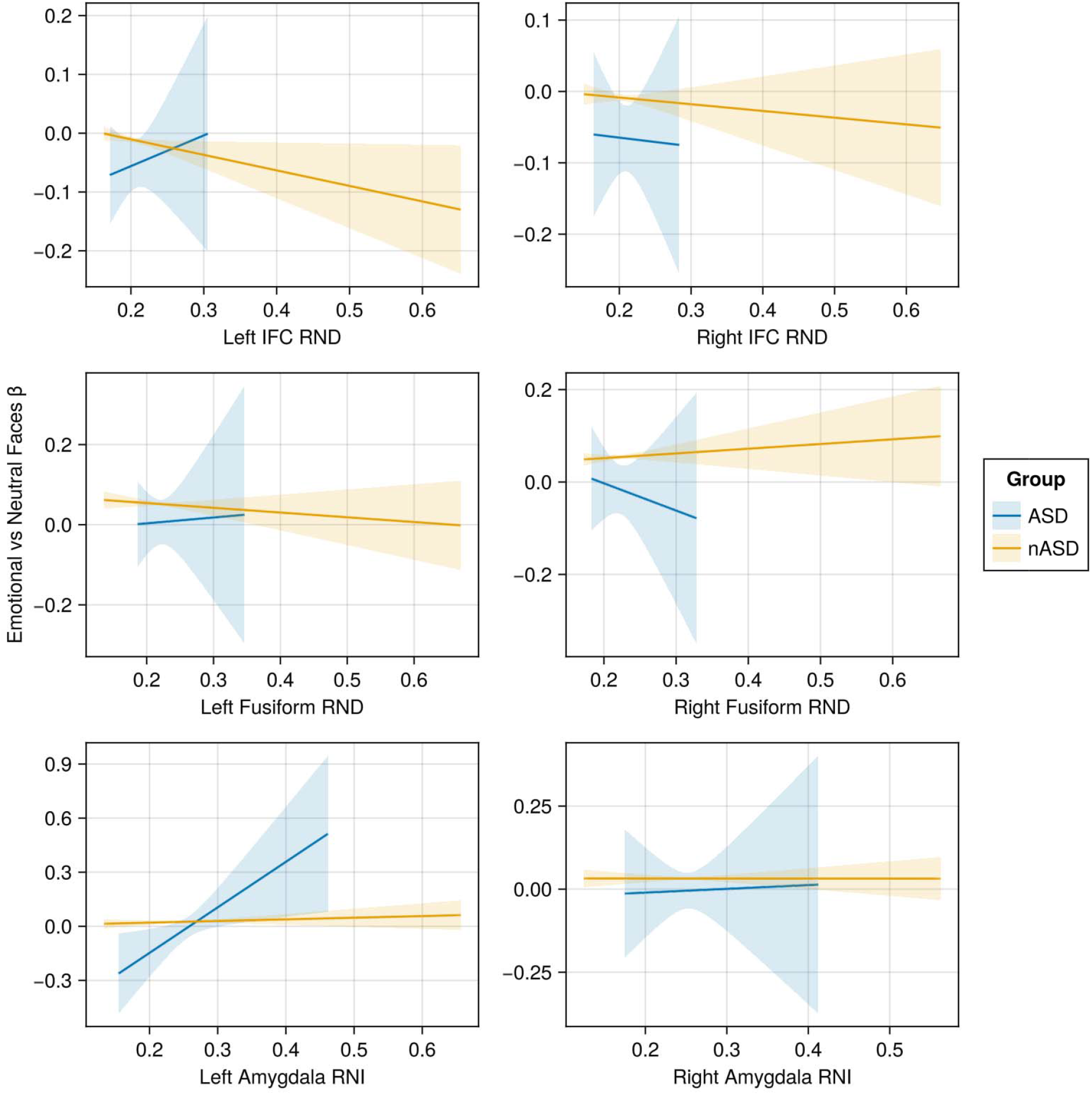
Along the y-axis are the beta values while processing emotional faces contrasted with the beta weights while processing neutral faces, representing the functional activity uniquely attributed to processing emotional expression in faces. Along the x-axis are measures of neuron density for each region. Blue and yellow lines indicate the linear model between these axes for the ASD and nASD groups, respectively. These relationships represent the raw linear relationship prior to inclusion of hierarchical control variables in the linear model reported in the results. **IFC**: The ASD group had an increase in RND associated with an increase in functional activity related to emotional processing of faces in the left IFC, whereas the nASD group had a negative association between RND and functional activity. Both groups had a decrease in functional activity related to RND in the right IFC. **Fusiform**: There was a modest negative association between RND and functional activity in the left fusiform for the nASD group and a modest positive association for the ASD group. There was a modest positive relationship between RND and functional activity in the right fusiform for the nASD group and a negative relationship between the ASD group. **Amygdala**: There was a positive relationship between RNI and functional activity in the amygdala for both groups. However, this relationship was only modest in the nASD group with a greater correlation in the ASD group, particularly in in the left amygdala.

Similar to those findings in the left IFC, the left fusiform initially demonstrated a negative association in the nASD group and a positive association in the ASD group (see Figure 1 middle-left). However, once controlling for confounding variables there was only a modest positive correlation in the nASD group (β=0.0327871, SE=0.490262, p-value=0.9467) and the ASD group (β=0.12821, SE=0.487922, p-value=0.7927). In contrast, the right fusiform had a modest negative association between RND and functional activity in the nASD group (β=−0.226399, SE=0.514337, p-value=0.6598) and ASD group (β=−0.337724, SE=0.514183, p-value=0.5113).

RNI was used to assess density of neuron cell bodies at the amygdala. Greater functional activity was elicited while processing emotional faces compared to neutral faces by those with an increased density of neuron cell bodies in the amygdala bilaterally (left: β=1.26596, SE=0.413497, p-value=0.0022; right: β=0.055502, SE=0.506337, p-value=0.9127). Although the positive association was consistent in the ASD group, they demonstrated a greater positive association in the left amygdala (left: β=1.23408, SE=0.412041, p-value=0.0027; right: β=0.0524766, SE=0.506349, p-value=0.9175).

## 4 Discussion

To gain insight into the historically inconsistent functional imaging findings related to the processing of emotional faces in ASD, the present study incorporated recently identified cytoarchitectural measures that differed in those with ASD. The use of imaging from the ABCD study (Barch et al., 2018, 2021; Casey et al., 2018; Hagler et al., 2019), a previously identified population of individuals with ASD, and relevant cytoarchitectural measures (Christensen et al., 2024) allowed for a more focused investigation than previously possible. Specifically, the relationship between RNI in the amygdala, RND in the fusiform, and RND in the IFC was evaluated in the context of processing emotional faces during fMRI scanning. Although these regions have been found to differ functionally in ASD in previous studies (Monk et al., 2010; Rämä & Courtney, 2005), the present study did not identify significant findings in the fusiform or IFC. However, a significant reduction in functional activity in the left amygdala while processing emotional faces in ASD was found. Furthermore, the ASD group were found to differ in the relationship between functional activity attributable to processing emotional faces and cytoarchitecture in the left amygdala, potentially demonstrating a differential dependence on cytoarchitecture in ASD for socio-emotional functioning.

The significant reduction in functional activity of the left amygdala in the ASD group when processing emotional faces is inconsistent with prior research where increased functional activity was found in the amygdala while processing emotional faces (Monk et al., 2010). The current design had the benefit of collecting imaging data across four timepoints spanning a 6-year period, with multiple trials of the EN-Back task. The significant reduction in functional activity identified in the ASD group potentially represents a more robust characterization of decreased mean functional activity than previous studies. However, functional activity is also dependent on temporal factors such as habituation (Kleinhans et al., 2016; Tam et al., 2017) or functional connectivity (Guo et al., 2016). Therefore, additional characterization of functional activity during processing of emotional faces in ASD is necessary to fully elucidate this relationship.

In addition to mean functional activity, the relationship between emotional recognition and the amygdala may be related to changes in structural connectivity throughout development (Saygin et al., 2015). The present study found a significant correlation between functional activity while processing emotional faces and estimated neuron cell body density in the left amygdala (see Figure 1 left-bottom). In addition to the overall positive association, the ASD group had an even greater positive correlation. The amygdala is of interest in ASD as other studies have posited that ASD is characterized by periods of premature maturation of the amygdala (Groen et al., 2010; Nordahl et al., 2012). Although the present study included statistical modelling to control for differences across timepoints, a differential trajectory in amygdalar development between groups may potentially confound findings. Interestingly, post-hoc analysis demonstrated a strong positive correlation between timepoint and amygdalar neuron cell body density bilaterally, with non-significant associations in other regions (see Figure 2). However, this relationship was consistent across both the ASD and nASD groups, suggesting neurodevelopmental changes in the amygdala from ages 9-16 represent a common process in both populations. Therefore, the significant relationship between functional activity while processing of emotional faces and RNI in the left amygdala is likely independent of unique developmental changes in the amygdala across this developmental timespan.

**Figure 2.**
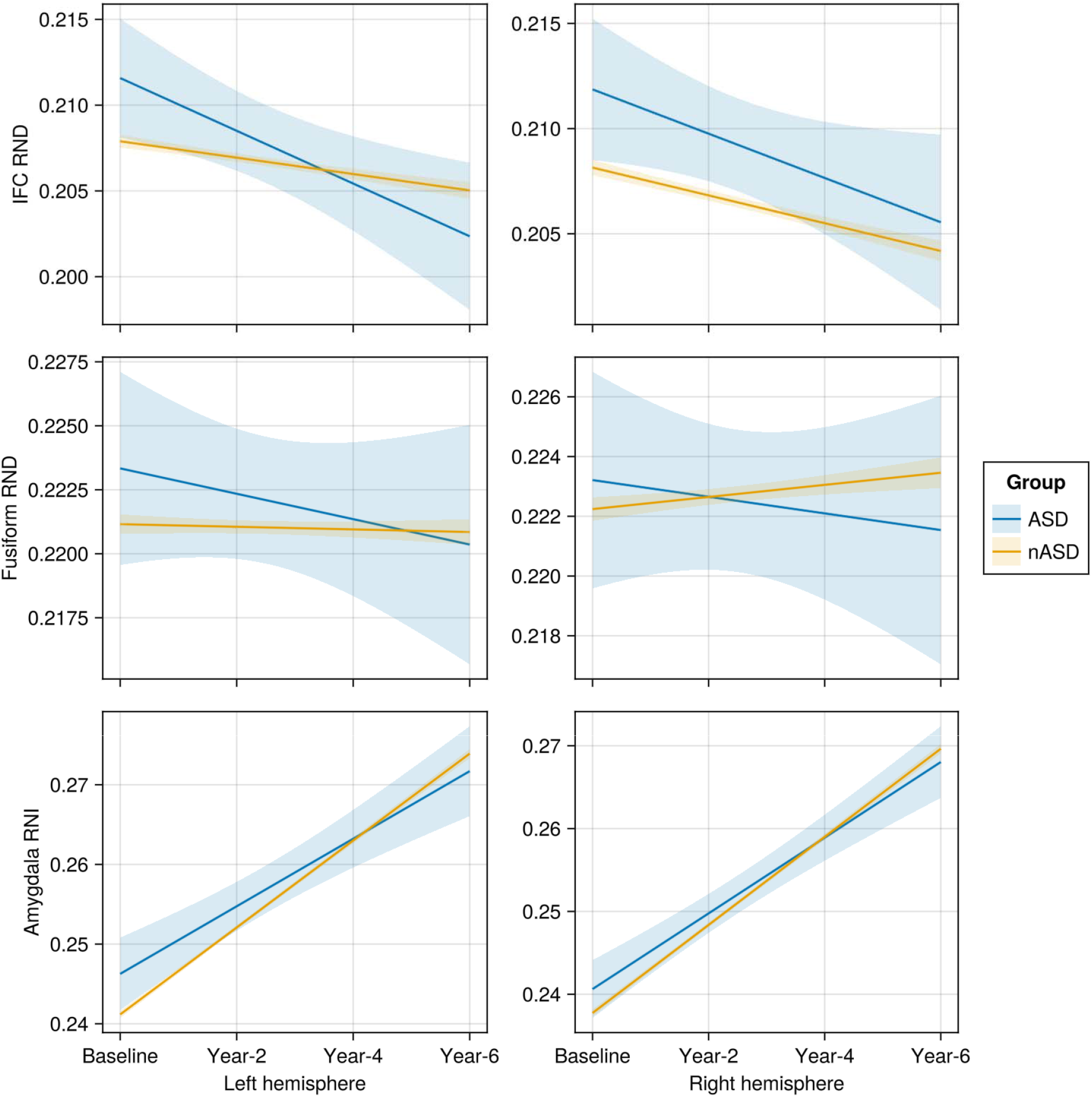
The linear relationship between neuron density measures (y-axis) and each timepoint in order (x-axis). **IFC**: Both the ASD group and nASD group had a modest reduction in RND bilaterally in the IFC across the 6-year timespan. **Fusiform**: The ASD group demonstrated a similarly modest reduction in in RND in the fusiform while the nASD group demonstrated a modest positive increase in fusiform RND. **Amygdala**: Both the ASD and nASD group had a significant increase in amygdalar RNI bilaterally.

There were no significant findings specific to the fusiform in the ASD group. This is inconsistent with prior research that found individuals with ASD demonstrated reduced functional activity in the fusiform when processing faces (Nickl-Jockschat et al., 2015). However, a significant positive correlation between RND and functional activation in the right fusiform for the nASD group was found. The absence of a similar relationship in the ASD group may be due to an overall shift toward lower RND values (see Figure 1 row 2), requiring a greater sample size to compensate for the absence of higher values driving a correlation. Alternatively, the complex relationship between the fusiform and other structures involved in processing emotional faces may require a more nuanced model informed by more modalities (Floris et al., 2025).

Prior literature has identified decreased functional activity in the IFC of individuals with ASD when performing tasks that required interpretation of emotional faces (Kilroy et al., 2021) or working memory of emotional faces (Rämä & Courtney, 2005). Although the present study identified decreased functional activity in the IFC of those in the ASD group when processing emotional faces, these findings were not significant when accounting for additional confounding variables. The current study sought to characterize the relationship between each region individually. However, recent analyses have demonstrated that the IFC may play an important role in mediating function of brain regions, particularly in relationship to mental health (Q. Li et al., 2025). Therefore, it is important that future s model the relationship between regions of the brain when processing emotional faces.

A meaningful difference in functional activity based on sex was not observed in the present study. These findings may be influenced by the small number of females in the ASD group (less than 20 females identified, see Table 1). This overrepresentation of males is consistent with the sex ratios observed in ASD across the global population (Elsabbagh et al., 2012). However, males with ASD have been found to have greater connectivity between functionally homotopic voxels in the left fusiform and right inferior frontal gyrus, when compared to females with ASD (H. Li et al., 2024). Sex-specific differences in brain structure in ASD have also been identified (e.g., brain symmetry (Deng & Wang, 2021), frontal and superior temporal cortices (Retico et al., 2016)). Although, sex-specific differences within the ASD population are frequently observed, the typical course of early neurodevelopment in nASD has been found to differ between males and females (Gilmore et al., 2007), with some of these typical development differences also identified in ASD (Retico et al., 2016). Therefore, further investigation accounting for demographic differences as well as less frequently utilized measures of the brain (e.g., symmetry, homotopic activity) are necessary to confirm that there are no sex-based differences in functional activity when processing emotional faces.

Given the novel investigation of in vivo estimates of neuron density and its association with functional activation during the processing of emotional faces, there are some important limitations to be considered. The current study only considers individuals between 9 and 16 years of age. Although narrowing the age-range avoids obscuring findings with changes unique to specific neurodevelopmental stages in ASD (Khundrakpam et al., 2017), it limits generalizability of findings. Furthermore, statistical models included control variables previously identified in the ABCD study (Barch et al., 2018; Christensen et al., 2021; Hagler et al., 2019), but were unable to explicitly account for comorbidities common to ASD or neuropsychiatric disorders superimposed on ASD. Therefore, future investigation should further explore these findings across a greater span of development while monitoring their stability in cohorts with known comorbidities.

Controlling for environmental variables that can potentially influence neurodevelopment has become an increasing point of concern in research. The effect of environmental variables across time has previously been considered specifically in the ABCD population (Gonzalez et al., 2021), with additional recommendations for management across the six year timespan (Hawes et al., 2025). The present study attempted to consider these factors in both data management and modeling. However, there remains additional complexity when incorporating relatively novel biomarkers, such as those characterizing in vivo neuron density. As stated previously, this was briefly explored to ensure validity of findings related to mean outcome measures across the 6-year span. However, variable neurodevelopment is of particular concern in ASD (Groen et al., 2010; Nordahl et al., 2012). Therefore, future research will need to continue identifying differential neurodevelopment in ASD but also the effects of environmental exposures on its trajectory.

## 5 Conclusion

In summary, the present study evaluated functional activity while processing emotional faces while controlling for processing neutral faces, finding a relative decrease in functional activity in the left amygdala of the ASD group compared to the nASD group. An additional interaction between ASD group and RNI was found, demonstrating a unique positive correlation between cell body density in the left amygdala and functional activity in the ASD group. In contrast, the IFC and fusiform had no significant findings between functional activity and the ASD group. Although the lack of findings in other regions is inconsistent with prior investigations, the extent to which these findings may differ in a more heterogeneous representation of ASD is still unknown. Finally, further investigation should be pursued to determine how functional activity in the left amygdala is related to the unique development of cell bodies in ASD and its potential application to clinical interventions.

## Acknowledgements

Data used in the preparation of this article were obtained from the Adolescent Brain Cognitive Development^SM^ (ABCD) Study (https://abcdstudy.org), held in the NIMH Data Archive (NDA). This is a multisite, longitudinal study designed to recruit more than 10,000 children age 9–10 and follow them over 10 years into early adulthood. Special thanks to the ABCD research team at the Cognitive Neurophysiology Laboratory (CNL) at the University of Rochester for their tireless work with the children and adolescents of the ABCD cohort.

## Funding

The ABCD Study® is supported by the National Institutes of Health and additional federal partners under award numbers U01DA041048, U01DA050989, U01DA051016, U01DA041022, U01DA051018, U01DA051037, U01DA050987, U01DA041174, U01DA041106, U01DA041117, U01DA041028, U01DA041134, U01DA050988, U01DA051039, U01DA041156, U01DA041025, U01DA041120, U01DA051038, U01DA041148, U01DA041093, U01DA041089, U24DA041123, U24DA041147. Neuroimaging at the University of Rochester ABCD site is conducted through the Translational Neuroimaging and Neurophysiology Core of the University of Rochester Intellectual and Developmental Disabilities Research Center (UR-IDDRC) which is supported by a center grant from the Eunice Kennedy Shriver National Institute of Child Health and Human Development (P50 HD103536 – to JJF). A full list of supporters is available at https://abcdstudy.org/federal-partners.html. A listing of participating sites and a complete listing of the study investigators can be found at https://abcdstudy.org/consortium_members/. ABCD consortium investigators designed and implemented the study and/or provided data but did not necessarily participate in the analysis or writing of this report. This manuscript reflects the views of the authors and may not reflect the opinions or views of the NIH or ABCD consortium investigators. ZPC’s work on this project was supported by the University of Rochester CTSA award number TL1 TR002000 from the National Center for Advancing Translational Sciences of the National Institutes of Health. Adolescent Brain and Cognitive Development study is a service mark of the U.S. Department of Health and Human Services.

## Data Sharing

The ABCD data repository grows and changes over time. The ABCD data used in this report came from DOI: 10.15154/z563-zd24. Code used in analysis is available by request from the authors.

## Conflict-of-Interest Statement

The authors have no financial interests that would present a conflict-of-interest in relation to the work reported herein.

## List of Abbreviations

(ASD): autism spectrum diagnosis/disorder
(nASD): no reported diagnosis of ASD
(EN-Back): emotional n-back
(fMRI): function magnetic resonance imaging
(MRI): magnetic resonance imaging
(DWI): diffusion weighted imaging
(IFC): inferior frontal cortex
(RSI): restriction spectrum imaging
(RND): restricted normalized direction diffusion
(RNI): restricted normalized isotropic diffusion

## Author Contributions

ZPC performed the primary data analyses and statistical tests, created the illustrations, and wrote the first substantial draft of the manuscript. JJF and EGF are site PIs for the ABCD consortium at the University of Rochester and coordinated data collection and project management. ZPC, JJF and EGF collectively conceived of the study and consulted regularly during the development of the analyses and data representations. JJF and EGF provided editorial input to ZPC during multiple drafts of the manuscript. All authors approve of this final version for publication, attest to the accuracy of the work reported, and agree to be fully accountable for all aspects of the work.

## Notes

### Competing Interest Statement

The authors have declared no competing interest.

## References

American Psychiatric Association (Ed.). (2013). Diagnostic and statistical manual of mental disorders: DSM-5 (5th ed). American Psychiatric Association.

Barch, D. M., Albaugh, M. D., Avenevoli, S., Chang, L., Clark, D. B., Glantz, M. D., Hudziak, J. J., Jernigan, T. L., Tapert, S. F., Yurgelun-Todd, D., Alia-Klein, N., Potter, A. S., Paulus, M. P., Prouty, D., Zucker, R. A., & Sher, K. J. (2018). Demographic, physical and mental health assessments in the adolescent brain and cognitive development study: Rationale and description. Developmental Cognitive Neuroscience, 32, 55–66. 10.1016/j.dcn.2017.10.010

Barch, D. M., Albaugh, M. D., Baskin-Sommers, A., Bryant, B. E., Clark, D. B., Dick, A. S., Feczko, E., Foxe, J. J., Gee, D. G., Giedd, J., Glantz, M. D., Hudziak, J. J., Karcher, N. R., LeBlanc, K., Maddox, M., McGlade, E. C., Mulford, C., Nagel, B. J., Neigh, G., … Xie, L. (2021). Demographic and mental health assessments in the adolescent brain and cognitive development study: Updates and age-related trajectories. Developmental Cognitive Neuroscience, 52, 101031. 10.1016/j.dcn.2021.101031

Barch, D. M., Burgess, G. C., Harms, M. P., Petersen, S. E., Schlaggar, B. L., Corbetta, M., Glasser, M. F., Curtiss, S., Dixit, S., Feldt, C., Nolan, D., Bryant, E., Hartley, T., Footer, O., Bjork, J. M., Poldrack, R., Smith, S., Johansen-Berg, H., Snyder, A. Z., & Van Essen, D. C. (2013). Function in the human connectome: Task-fMRI and individual differences in behavior. NeuroImage, 80, 169–189. 10.1016/j.neuroimage.2013.05.033

Bates, D., Mächler, M., Bolker, B., & Walker, S. (2014). Fitting Linear Mixed-Effects Models using lme4. 1406.5823 [Stat]. http://arxiv.org/abs/1406.5823

Benjamini, Y., & Hochberg, Y. (1995). Controlling the False Discovery Rate: A Practical and Powerful Approach to Multiple Testing. Journal of the Royal Statistical Society: Series B (Methodological), 57(1), 289–300. 10.1111/j.2517-6161.1995.tb02031.x

Buxhoeveden, D. P., Semendeferi, K., Buckwalter, J., Schenker, N., Switzer, R., & Courchesne, E. (2006). Reduced minicolumns in the frontal cortex of patients with autism. Neuropathology and Applied Neurobiology, 32(5), 483–491. 10.1111/j.1365-2990.2006.00745.x

Casanova, M. F., Buxhoeveden, D. P., Switala, A. E., & Roy, E. (2002). Minicolumnar pathology in autism. Neurology, 58(3), 428–432. 10.1212/WNL.58.3.428

Casanova, M., & Trippe, J. (2009). Radial cytoarchitecture and patterns of cortical connectivity in autism. Philosophical Transactions of the Royal Society B: Biological Sciences, 364(1522), 1433–1436. 10.1098/rstb.2008.0331

Casey, B. J., Cannonier, T., Conley, M. I., Cohen, A. O., Barch, D. M., Heitzeg, M. M., Soules, M. E., Teslovich, T., Dellarco, D. V., Garavan, H., Orr, C. A., Wager, T. D., Banich, M. T., Speer, N. K., Sutherland, M. T., Riedel, M. C., Dick, A. S., Bjork, J. M., Thomas, K. M., … Dale, A. M. (2018). The Adolescent Brain Cognitive Development (ABCD) study: Imaging acquisition across 21 sites. Developmental Cognitive Neuroscience, 32, 43–54. 10.1016/j.dcn.2018.03.001

Chaarani, B., Hahn, S., Allgaier, N., Adise, S., Owens, M. M., Juliano, A. C., Yuan, D. K., Loso, H., Ivanciu, A., Albaugh, M. D., Dumas, J., Mackey, S., Laurent, J., Ivanova, M., Hagler, D. J., Cornejo, M. D., Hatton, S., Agrawal, A., Aguinaldo, L., … the ABCD Consortium. (2021). Baseline brain function in the preadolescents of the ABCD Study. Nature Neuroscience, 24(8), 1176–1186. 10.1038/s41593-021-00867-9

Christensen, Z. P., Freedman, E. G., & Foxe, J. J. (2021). Caffeine exposure in utero is associated with structural brain alterations and deleterious neurocognitive outcomes in 9–10 year old children. Neuropharmacology, 186, 108479. 10.1016/j.neuropharm.2021.108479

Christensen, Z. P., Freedman, E. G., & Foxe, J. J. (2024). Autism is associated with in vivo changes in gray matter neurite architecture. Autism Research, aur.3239. 10.1002/aur.3239

Coad, B. M., Postans, M., Hodgetts, C. J., Muhlert, N., Graham, K. S., & Lawrence, A. D. (2020). Structural connections support emotional connections: Uncinate Fasciculus microstructure is related to the ability to decode facial emotion expressions. Neuropsychologia, 145, 106562. 10.1016/j.neuropsychologia.2017.11.006

Deng, Z., & Wang, S. (2021). Sex differentiation of brain structures in autism: Findings from a gray matter asymmetry study. Autism Research, aur.2506. 10.1002/aur.2506

Douglas Bates, Phillip Alday, Dave Kleinschmidt, José Bayoán Santiago Calderón, Andreas Noack, Tony Kelman, Milan Bouchet-Valat, Yakir Luc Gagnon, Simon Babayan, Patrick Kofod Mogensen, Morten Piibeleht, Michael Hatherly, Elliot Saba, & Antoine Baldassari. (2019). JuliaStats/MixedModels.jl: V2.2.0 [Computer software]. Zenodo. 10.5281/zenodo.3568326

Elbich, D. B., & Scherf, S. (2017). Beyond the FFA: Brain-behavior correspondences in face recognition abilities. NeuroImage, 147, 409–422. 10.1016/j.neuroimage.2016.12.042

Elsabbagh, M., Divan, G., Koh, Y.-J., Kim, Y. S., Kauchali, S., Marcín, C., Montiel-Nava, C., Patel, V., Paula, C. S., Wang, C., Yasamy, M. T., & Fombonne, E. (2012). Global Prevalence of Autism and Other Pervasive Developmental Disorders: Global epidemiology of autism. Autism Research, 5(3), 160–179. 10.1002/aur.239

Fatemi, S. H., Halt, A. R., Realmuto, G., Earle, J., Kist, D. A., Thuras, P., & Merz, A. (2002). Purkinje Cell Size Is Reduced in Cerebellum of Patients with Autism. Cellular and Molecular Neurobiology, 22(2), 171–175. 10.1023/A:1019861721160

Floris, D. L., Llera, A., Zabihi, M., Moessnang, C., Jones, E. J. H., Mason, L., Haartsen, R., Holz, N. E., Mei, T., Elleaume, C., Vieira, B. H., Pretzsch, C. M., Forde, N. J., Baumeister, S., Dell’Acqua, F., Durston, S., Banaschewski, T., Ecker, C., Holt, R. J., … Langer, N. (2025). A multimodal neural signature of face processing in autism within the fusiform gyrus. Nature Mental Health, 3(1), 31–45. 10.1038/s44220-024-00349-4

Garavan, H., Bartsch, H., Conway, K., Decastro, A., Goldstein, R. Z., Heeringa, S., Jernigan, T., Potter, A., Thompson, W., & Zahs, D. (2018). Recruiting the ABCD sample: Design considerations and procedures. Developmental Cognitive Neuroscience, 32, 16–22. 10.1016/j.dcn.2018.04.004

Gilmore, J. H., Lin, W., Corouge, I., Vetsa, Y. S. K., Smith, J. K., Kang, C., Gu, H., Hamer, R. M., Lieberman, J. A., & Gerig, G. (2007). Early Postnatal Development of Corpus Callosum and Corticospinal White Matter Assessed with Quantitative Tractography. American Journal of Neuroradiology, 28(9), 1789–1795. 10.3174/ajnr.A0751

Gonzalez, R., Thompson, E. L., Sanchez, M., Morris, A., Gonzalez, M. R., Feldstein Ewing, S. W., Mason, M. J., Arroyo, J., Howlett, K., Tapert, S. F., & Zucker, R. A. (2021). An update on the assessment of culture and environment in the ABCD Study®: Emerging literature and protocol updates over three measurement waves. Developmental Cognitive Neuroscience, 52, 101021. 10.1016/j.dcn.2021.101021

Griffin, J. W., Bauer, R., & Scherf, K. S. (2021). A quantitative meta-analysis of face recognition deficits in autism: 40 years of research. Psychological Bulletin, 147(3), 268–292. 10.1037/bul0000310

Groen, W., Teluij, M., Buitelaar, J., & Tendolkar, I. (2010). Amygdala and Hippocampus Enlargement During Adolescence in Autism. Journal of the American Academy of Child & Adolescent Psychiatry, 49(6), 552– 560. 10.1016/j.jaac.2009.12.023

Guo, X., Duan, X., Long, Z., Chen, H., Wang, Y., Zheng, J., Zhang, Y., Li, R., & Chen, H. (2016). Decreased amygdala functional connectivity in adolescents with autism: A resting-state fMRI study. Psychiatry Research: Neuroimaging, 257, 47–56. 10.1016/j.pscychresns.2016.10.005

Hagler, D. J., Hatton, SeanN., Cornejo, M. D., Makowski, C., Fair, D. A., Dick, A. S., Sutherland, M. T., Casey, B. J., Barch, D. M., Harms, M. P., Watts, R., Bjork, J. M., Garavan, H. P., Hilmer, L., Pung, C. J., Sicat, C. S., Kuperman, J., Bartsch, H., Xue, F., … Dale, A. M. (2019). Image processing and analysis methods for the Adolescent Brain Cognitive Development Study. NeuroImage, 116091. 10.1016/j.neuroimage.2019.116091

Hawes, S. W., Littlefield, A. K., Lopez, D. A., Sher, K. J., Thompson, E. L., Gonzalez, R., Aguinaldo, L., Adams, A. R., Bayat, M., Byrd, A. L., Castro-de-Araujo, L. F., Dick, A., Heeringa, S. F., Kaiver, C. M., Lehman, S. M., Li, L., Linkersdörfer, J., Maullin-Sapey, T. J., Neale, M. C., … Thompson, W. K. (2025). Longitudinal analysis of the ABCD® study. Developmental Cognitive Neuroscience, 72, 101518. 10.1016/j.dcn.2025.101518

Jernigan, T. L., Brown, S. A., Dale, A. M., Tapert, S. F., Sowell, E. R., Herting, M., Laird, A., Gonzalez, R., Squeglia, L., Gray, K., Paulus, M. P., Aupperle, R., Feldstein Ewing, S. W., Nagel, B. J., Fair, D. A., Baker, F., Müller Oehring, E., Bookheimer, S. Y., Dapretto, M., … Gee, D. (2025). ABCD Study(R) Data Release 6.0 [Dataset]. Lasso Informatics US Inc. 10.82525/JY7N-G441

Kanwisher, N., & Barton, J. J. S. (2011). The Functional Architecture of the Face System: Integrating Evidence from fMRI and Patient Studies. Oxford University Press. 10.1093/oxfordhb/9780199559053.013.0007

Kanwisher, N., McDermott, J., & Chun, M. M. (1997). The Fusiform Face Area: A Module in Human Extrastriate Cortex Specialized for Face Perception. The Journal of Neuroscience, 17(11), 4302–4311. 10.1523/JNEUROSCI.17-11-04302.1997

Kemper, T. L., & Bauman, M. (1998). Neuropathology of Infantile Autism: Journal of Neuropathology and Experimental Neurology, 57(7), 645–652. 10.1097/00005072-199807000-00001

Khundrakpam, B. S., Lewis, J. D., Kostopoulos, P., Carbonell, F., & Evans, A. C. (2017). Cortical Thickness Abnormalities in Autism Spectrum Disorders Through Late Childhood, Adolescence, and Adulthood: A Large-Scale MRI Study. Cerebral Cortex, 27(3), 1721–1731. 10.1093/cercor/bhx038

Kietzmann, T. C., Poltoratski, S., König, P., Blake, R., Tong, F., & Ling, S. (2015). The Occipital Face Area Is Causally Involved in Facial Viewpoint Perception. The Journal of Neuroscience: The Official Journal of the Society for Neuroscience, 35(50), 16398–16403. 10.1523/JNEUROSCI.2493-15.2015

Kilroy, E., Harrison, L., Butera, C., Jayashankar, A., Cermak, S., Kaplan, J., Williams, M., Haranin, E., Bookheimer, S., Dapretto, M., & Aziz-Zadeh, L. (2021). Unique deficit in embodied simulation in autism: An FMRI study comparing autism and developmental coordination disorder. Human Brain Mapping, 42(5), 1532–1546. 10.1002/hbm.25312

Klein, A., & Tourville, J. (2012). 101 Labeled Brain Images and a Consistent Human Cortical Labeling Protocol. Frontiers in Neuroscience, 6. 10.3389/fnins.2012.00171

Kleinhans, N. M., Richards, T., Greenson, J., Dawson, G., & Aylward, E. (2016). Altered Dynamics of the fMRI Response to Faces in Individuals with Autism. Journal of Autism and Developmental Disorders, 46(1), 232–241. 10.1007/s10803-015-2565-8

Kleinhans, N. M., Richards, T., Johnson, L. C., Weaver, K. E., Greenson, J., Dawson, G., & Aylward, E. (2011). fMRI evidence of neural abnormalities in the subcortical face processing system in ASD. NeuroImage, 54(1), 697–704. 10.1016/j.neuroimage.2010.07.037

Li, H., Zhang, Q., Duan, T., Li, J., Shi, L., Hua, Q., Li, D., Ji, G.-J., Wang, K., & Zhu, C. (2024). Sex differences in brain functional specialization and interhemispheric cooperation among children with autism spectrum disorders. Scientific Reports, 14(1), 22096. 10.1038/s41598-024-72339-6

Li, Q., Cao, M., Stein, D. J., Sahakian, B. J., Jia, T., Langley, C., Gu, Z., Hou, W., Lu, H., Cao, L., Lin, J., Shi, R., Banaschewski, T., Bokde, A. L. W., Desrivières, S., Flor, H., Grigis, A., Garavan, H., Gowland, P., … IMAGEN Consortium. (2025). Cognitive predictors of mental health trajectories are mediated by inferior frontal and occipital development during adolescence. Molecular Psychiatry, 30(7), 3055–3068. 10.1038/s41380-025-02912-6

McKechanie, A. G., Lawrie, S. M., Whalley, H. C., & Stanfield, A. C. (2022). A functional MRI facial emotion-processing study of autism in individuals with special educational needs. Psychiatry Research: Neuroimaging, 320, 111426. 10.1016/j.pscychresns.2021.111426

Monk, C. S., Weng, S.-J., Wiggins, J. L., Kurapati, N., Louro, H. M. C., Carrasco, M., Maslowsky, J., Risi, S., & Lord, C. (2010). Neural circuitry of emotional face processing in autism spectrum disorders. Journal of Psychiatry and Neuroscience, 35(2), 105–114. 10.1503/jpn.090085

Moul, C., Cauchi, A., Hawes, D. J., Brennan, J., & Dadds, M. R. (2015). Differentiating Autism Spectrum Disorder and Overlapping Psychopathology with a Brief Version of the Social Responsiveness Scale. Child Psychiatry & Human Development, 46(1), 108–117. 10.1007/s10578-014-0456-4

Nickl-Jockschat, T., Rottschy, C., Thommes, J., Schneider, F., Laird, A. R., Fox, P. T., & Eickhoff, S. B. (2015). Neural networks related to dysfunctional face processing in autism spectrum disorder. Brain Structure & Function, 220(4), 2355–2371. 10.1007/s00429-014-0791-z

Nordahl, C. W., Scholz, R., Yang, X., Buonocore, M. H., Simon, T., Rogers, S., & Amaral, D. G. (2012). Increased rate of amygdala growth in children aged 2 to 4 years with autism spectrum disorders: A longitudinal study. Archives of General Psychiatry, 69(1), 53–61. 10.1001/archgenpsychiatry.2011.145

Pedersen, M. L., Alnæs, D., Van Der Meer, D., Fernandez-Cabello, S., Berthet, P., Dahl, A., Kjelkenes, R., Schwarz, E., Thompson, W. K., Barch, D. M., Andreassen, O. A., & Westlye, L. T. (2023). Computational Modeling of the n-Back Task in the ABCD Study: Associations of Drift Diffusion Model Parameters to Polygenic Scores of Mental Disorders and Cardiometabolic Diseases. Biological Psychiatry: Cognitive Neuroscience and Neuroimaging, 8(3), 290–299. 10.1016/j.bpsc.2022.03.012

Rämä, P., & Courtney, S. M. (2005). Functional topography of working memory for face or voice identity. NeuroImage, 24(1), 224–234. 10.1016/j.neuroimage.2004.08.024

Retico, A., Giuliano, A., Tancredi, R., Cosenza, A., Apicella, F., Narzisi, A., Biagi, L., Tosetti, M., Muratori, F., & Calderoni, S. (2016). The effect of gender on the neuroanatomy of children with autism spectrum disorders: A support vector machine case-control study. Molecular Autism, 7(1), 5. 10.1186/s13229-015-0067-3

Sato, W., Usui, N., Sawada, R., Kondo, A., Toichi, M., & Inoue, Y. (2021). Impairment of emotional expression detection after unilateral medial temporal structure resection. Scientific Reports, 11(1), 20617. 10.1038/s41598-021-99945-y

Saygin, Z. M., Osher, D. E., Koldewyn, K., Martin, R. E., Finn, A., Saxe, R., Gabrieli, J. D. E., & Sheridan, M. (2015). Structural connectivity of the developing human amygdala. PloS One, 10(4), e0125170. 10.1371/journal.pone.0125170

Schumann, C. M., & Amaral, D. G. (2006). Stereological Analysis of Amygdala Neuron Number in Autism. Journal of Neuroscience, 26(29), 7674–7679. 10.1523/JNEUROSCI.1285-06.2006

Solomon-Harris, L. M., Mullin, C. R., & Steeves, J. K. E. (2013). TMS to the “occipital face area” affects recognition but not categorization of faces. Brain and Cognition, 83(3), 245–251. 10.1016/j.bandc.2013.08.007

Stoner, R., Chow, M. L., Boyle, M. P., Sunkin, S. M., Mouton, P. R., Roy, S., Wynshaw-Boris, A., Colamarino, S. A., Lein, E. S., & Courchesne, E. (2014). Patches of Disorganization in the Neocortex of Children with Autism. New England Journal of Medicine, 370(13), 1209–1219. 10.1056/NEJMoa1307491

Tam, F. I., King, J. A., Geisler, D., Korb, F. M., Sareng, J., Ritschel, F., Steding, J., Albertowski, K. U., Roessner, V., & Ehrlich, S. (2017). Altered behavioral and amygdala habituation in high-functioning adults with autism spectrum disorder: An fMRI study. Scientific Reports, 7(1), 13611. 10.1038/s41598-017-14097-2

Uljarevic, M., & Hamilton, A. (2013). Recognition of Emotions in Autism: A Formal Meta-Analysis. Journal of Autism and Developmental Disorders, 43(7), 1517–1526. 10.1007/s10803-012-1695-5

Vanneau, T. (2025). Neural oscillatory dynamics reveal altered top-down and integrative mechanisms during face processing in autistic children and unaffected siblings of autistic children.

Vuilleumier, P., Richardson, M. P., Armony, J. L., Driver, J., & Dolan, R. J. (2004). Distant influences of amygdala lesion on visual cortical activation during emotional face processing. Nature Neuroscience, 7(11), 1271– 1278. 10.1038/nn1341

Webb, S. J., Faja, S., & Dawson, G. (2011). Face Processing in Autism. Oxford University Press. 10.1093/oxfordhb/9780199559053.013.0043

White, N. S., Leergaard, T. B., D’Arceuil, H., Bjaalie, J. G., & Dale, A. M. (2013). Probing tissue microstructure with restriction spectrum imaging: Histological and theoretical validation. Human Brain Mapping, 34(2), 327–346. 10.1002/hbm.21454

Wong, N. M., Dipasquale, O., Turkheimer, F., Findon, J. L., Wichers, R. H., Dimitrov, M., Murphy, C. M., Stoencheva, V., Robertson, D. M., Murphy, D. G., Daly, E., & McAlonan, G. M. (2022). Differences in social brain function in autism spectrum disorder are linked to the serotonin transporter: A randomised placebo-controlled single-dose crossover trial. Journal of Psychopharmacology, 36(6), 723–731. 10.1177/02698811221092509

